# Enterohaemorrhagic *E. coli* utilizes an AND-OR logic gate to regulate expression of an outer membrane haem receptor

**DOI:** 10.1101/2021.08.24.457593

**Authors:** Brandon M. Sy, Jai J. Tree

## Abstract

To sense the transition from environment to host, bacteria use a range of environmental cues to control expression of virulence genes. Iron is tightly sequestered in host tissues and in the human pathogen enterohaemorrhagic *E. coli* (EHEC) iron-limitation induces transcription of the outermembrane haem transporter encoded by *chuAS*. ChuA expression is post-transcriptionally activated at 37°C by a FourU RNA thermometer ensuring that the haem receptor is only expressed under low iron, high temperature conditions that indicate the host. Here we demonstrate that expression of *chuA* is also independently regulated by the cAMP-responsive sRNA CyaR and transcriptional terminator Rho. These results indicate that *chuAS* expression is regulated at the transcription initiation, transcript elongation, and translational level. The natural dependence of these processes creates a hierarchy of regulatory AND and OR logic gates that integrate information about the local environment. We show that the logic of the *chuA* regulatory circuit is activated under conditions that satisfy low iron AND (low glucose OR high temperature). We speculate that additional sensing of a gluconeogenic environment allows further precision in determining when EHEC is at the gastrointestinal epithelium of the host. With previous studies, it appears that the *chuAS* transcript is controlled by eight regulatory inputs that control expression through six different transcriptional and post-transcriptional mechanisms. The results highlight the ability of regulatory sRNAs to integrate multiple environmental signals into a conditional hierarchy of signal input.

## INTRODUCTION

During infection, nutrients required by pathogenic bacteria are withheld by the host in a process known as nutritional immunity [1,2]. An example can be found in iron, a transition metal that is essential for pathogenic bacteria due to its role in essential physiological processes, such as respiration [3,4]. To deprive pathogens of this vital nutrient, the host sequesters iron in the high affinity iron binding molecules haem, ferritin and transferrin [5,6]. These ensure that the concentration of extracellular free iron in the host is too low to support bacterial growth and infection. In order to overcome this nutritional deficit, pathogens such as *H. influenzae, C. jejuni, P. aeruginosa* and uropathogenic *E. coli* have evolved multiple systems that allow for the scavenging of iron, these include siderophores and haem uptake systems [4,7–15]

Enterohaemorrhagic *Escherichia coli* (EHEC) is an enteric pathogen that causes haemorrhagic colitis which can progress into potentially fatal haemolytic uremic syndrome. The primary reservoir of this pathogen are ruminants and transmission mainly occurs via the consumption of contaminated foodstuffs [16–18]. EHEC can express an outer membrane haem receptor from the iron regulated *chu* haem uptake operon [19]. The *chu* locus is homologous to the *shu* locus found in *Shigella*, and contains genes that allow for haem uptake, transport, utilization and degradation. The locus consists of a bicistronic operon (*chuAS)* as well as two polycistronic operons (*chuTWXY* and *chuUV*). ChuA is a TonB-dependent outer membrane haem receptor that imports haem into the periplasm [20]. The periplasmic binding protein ChuT binds to haem, and via the ABC transporter ChuUV, is shuttled into the cytoplasm. Haem is processed by the haem oxygenase ChuS that converts haem into biliverdin, carbon monoxide and free iron. This breakdown also prevents haem toxicity [21]. Under anaerobic conditions, this role is taken up by the SAM-methyltransferase ChuW and the anaerobilin reductase ChuY [22,23].

The haem uptake operon is part of a larger regulatory network that allows EHEC to achieve iron homeostasis. Transcription of *chuA* is controlled by the iron-dependent transcriptional repressor Fur. In iron-rich conditions, Fur binds to iron which increases the affinity of Fur for its DNA-binding site ∼1000-fold [24]. Fur also regulates a Hfq-dependent small non-coding RNA (sRNA) RyhB that is central to maintaining iron homeostasis [25–28]. RyhB prevents the expression of non-essential proteins that require Fe as a cofactor such as the TCA cycle and respiration genes *sdhC* and *fumAC*, and iron storage genes such as bacterioferritin. Genes that contribute to iron uptake, such as the shikimate permease *shiA* and the colicin I receptor *cirA*, are positively regulated by this sRNA [27,29,30].

EHEC utilizes the Chu system when it is in an iron-poor, haem-rich host. For that reason, as well as the toxicity that accompanies haem and iron over-accumulation, it is necessary to precisely control expression of this operon. FourU RNA-thermometers are used in different bacterial pathogens, such as *Shigella dysenteriae, Yersinia pseudotuberculosis, Vibrio cholerae*, and *Salmonella typhimurium*, to regulate expression of their virulence factors in a temperature-dependent manner [31–34]. A FourU RNA-thermometer was found in the *chuA* 5’UTR that blocks the ribosomal binding site (RBS) and is formed at temperatures below 37°C, indicating that the pathogen is outside of the host [34]. This limits *<u>chuA</u>* expression outside of the host where haem is unlikely to be available.

Identification of binding sites for the small RNA chaperone Hfq in EHEC identified the phage-encoded sRNA AsxR that positively regulates the haem oxygenase *chuS* by sponging interactions with the negatively regulating sRNA, FnrS [35]. Further, analysis of the EHEC sRNA interactome revealed that the outer membrane haem receptor *chuA* was also negatively regulated by the EHEC-specific Hfq-binding sRNA Esr41 [36]. Here, we show that translation of *chuA* is activated by the Crp-cAMP regulated sRNA CyaR in a temperature- and Rho-termination-independent manner. Our results suggest that EHEC employs an AND-OR logic gate that integrates information on iron availability, temperature and carbon availability to sense its location within the host for appropriate expression of *chuA*.

## RESULTS

### The outer membrane haem receptor *chuA* is regulated by CyaR and ChiX

In previous work the binding sites of the small RNA chaperone Hfq were mapped using UV-crosslinking, denaturing purification and sequencing of Hfq-crosslinked RNA (termed CRAC) [35]. Deletions introduced at the site of protein-RNA crosslinking during reverse transcription of UV-crosslinked RNAs can be used to identify the site of direct Hfq-RNA contact. Analysis of our previous Hfq-CRAC dataset reveals that Hfq binds strongly to the 5’UTR of the outer membrane haem receptor *chuA* (Figure 1A & B). Canonically, gene regulation through sRNAs occurs through direct base-pairing of the sRNA to the ribosomal binding site (RBS) within the 5’UTR of mRNA targets. Hfq-CRAC sequencing reads and contact-dependant deletions peaked at the RBS consistent with RyhB and Esr41 negative regulation at this site [36,37]. A further Hfq-binding peak was located at position +160-+200 suggesting that *chuA* may be subject to additional regulatory inputs within the 5’UTR (Figure 1A and B).

**Figure 1:**
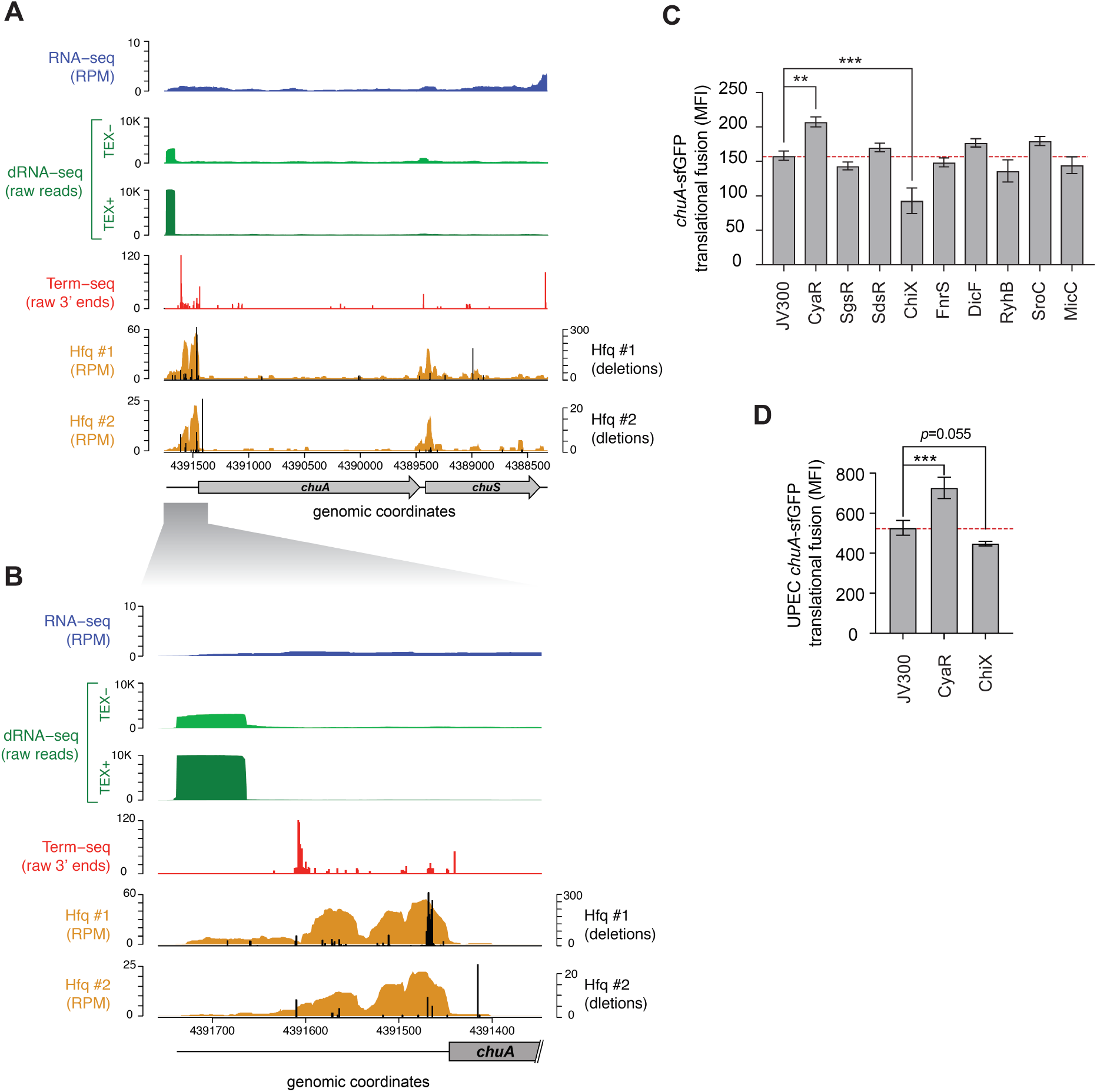
The *chuA* 5’UTR is regulated by Hfq-dependent small RNAs. **A**. RNA-sequencing data mapping to the *chuAS* operon. From top to bottom: total RNA-seq (dark blue; RPM), differential RNA-seq in samples untreated (light green) and treated (dark green) with TEX, Term-seq (red; raw 3’ end reads) (BioProject PRJNA601151), Hfq-binding sites from UV-crosslinking datasets (BioProject PRJNA197291). Sequencing deletions which indicate direct contact with Hfq are indicated in black. **B**. Same as A, but focusing on the 5’ UTR of *chuA*. **C**. Fluorescence measurements of the EHEC *chuA*-sfGFP fusion in the presence of predicted binding sRNAs. D. Fluorescence measurements of the UPEC *chuA*-sfGFP fusion in the presence of CyaR or ChiX. (*** *p <* 0.001, *\*\* p <* 0.01).

The *chuA* transcription start site is predicted to be ∼300 nt upstream of the *chuA* start codon [38]. To precisely map the transcription start site of *chuA* in EHEC str. Sakai, we analysed our previously published differential RNA-seq (dRNA-seq) data, which enriches for primary transcripts using terminator exonuclease (TEX) [39,40]. This showed that the +1 site for *chuA* is at position 4,391,736, and that the 5’UTR is 291 nucleotides long, which concurs with previous predictions (Figure 1B) [38].

In previous analyses we found that sRNA-mRNA seed sites are often positioned within Hfq-bound reads recovered by Hfq-CRAC, and that the Hfq binding site can be used to restrict the sequence space for predicting sRNA-mRNA interactions [35,41]. To identify sRNAs that may regulate *chuA*, IntaRNA [42,43] was used to predict interactions between the *chuA* 5’UTR and 44 known Hfq-dependent sRNAs [44]. Small RNAs that were predicted to interact within Hfq read peaks were retained. Through this analysis, we predicted that the sRNAs CyaR, SgrS, SdsR, ChiX, FnrS, RyhB, DicF, SroC and MicC may have regulatory effects on *chuA*.

To confirm these predicted interactions the 5’UTR of *chuA*, starting from the +1 site up until the 15^th^ codon of the coding sequence was fused to sfGFP by cloning into the plasmid pXG10SF [45]. The fluorescence of this translational fusion was monitored in *E. coli* strain DH5α in the presence and absence of the candidate sRNAs (Figure 1C). This revealed that the Hfq-dependent sRNAs CyaR and ChiX have an activating and repressive effect on *chuA*, respectively.

### CyaR and ChiX regulation of *chuA* is conserved in uropathogenic *E. coli*

In uropathogenic *E. coli* (UPEC), haem acquisition using *chuA* is required for maximal colonisation of the kidneys [11]. It has previously been shown that the 5’UTRs of EHEC and UPEC (ChuA_EHEC_ and ChuA_UPEC_) *chuA* are 85% similar [38]. The UPEC 5’UTR sequence is only 67.4% similar within the Hfq-binding site (versus 83.4% similarity for the 5’UTR as a whole) suggesting that *chuA*_UPEC_ may not be regulated by Hfq-dependant sRNAs. To test whether *chuA*_UPEC_ is post-transcriptionally regulated by CyaR and ChiX we constructed a translational fusion of *chuA*_UPEC_ with sfGFP and monitored fluorescence in the presence or absence of CyaR and ChiX. The *chuA*_UPEC_ fusion was activated by CyaR consistent with our results for *chuA*_EHEC_ but was not significantly repressed by ChiX (Figure 1D). Notably, the *chuA*_UPEC_ fusion was 55% more fluorescent than the *chuA*_EHEC_ fusion. This suggests that while *chuA* in both pathotypes is subject to regulation by CyaR, the divergent 5’UTR sequence has de-repressed *chuA* expression in UPEC.

### CyaR activates *chuA* translation through direct base-pairing

Both CyaR and ChiX are class II sRNAs that are known to modulate sRNA regulation by displacing sRNAs from Hfq when overexpressed [46,47]. To determine whether CyaR and ChiX control *chuA* through direct RNA-RNA interactions with the *chuA* 5’UTR or indirectly through titration of Hfq, point mutations were made in both *chuA* and the sRNAs CyaR and ChiX to disrupt the predicted interaction sites. A predicted 3 nt compensatory mutation was made in the top scoring *chuA*-ChiX interaction as predicted by IntaRNA. The M1 mutation made in ChiX resulted in the de-repression of *chuA*-GFP expression. However, this effect was not observed when the compensatory *chuA* M1 mutation was introduced. This suggested that while the nucleotides mutated in ChiX contributed to *chuA* regulation, they did not interact with the predicted *chuA* interaction site. A second set of compensatory point mutants (M2) were made for the next highest scoring interaction that used the same ChiX seed region that was previously tested. Mutations in either *chuA* or ChiX alone resulted in the disruption of the *chuA*-ChiX interaction, but testing the mutants together did not restore the repression of *chuA* (Supplementary Figure 1). These results indicated that while ChiX affects translation of *chuA*, it is not due to a direct interaction with the *chuA* M1 or M2 site and may occur indirectly.

CyaR is predicted to bind approximately 15 nt upstream of the *chuA* Shine-Dalgarno sequence. The interaction between CyaR and *chuA* results in a 1.3-fold activation of translation. Mutating CyaR at the M1 site reduced CyaR activation by a modest 8.1%, while the *chuA* point mutant completely abolished the interaction (Figure 2). Providing the CyaR-M1 mutant partially restore translational activation of *chuA*-M1, supporting a direct interaction between CyaR and the 5’UTR of *chuA* that activates translation.

**Figure 2:**
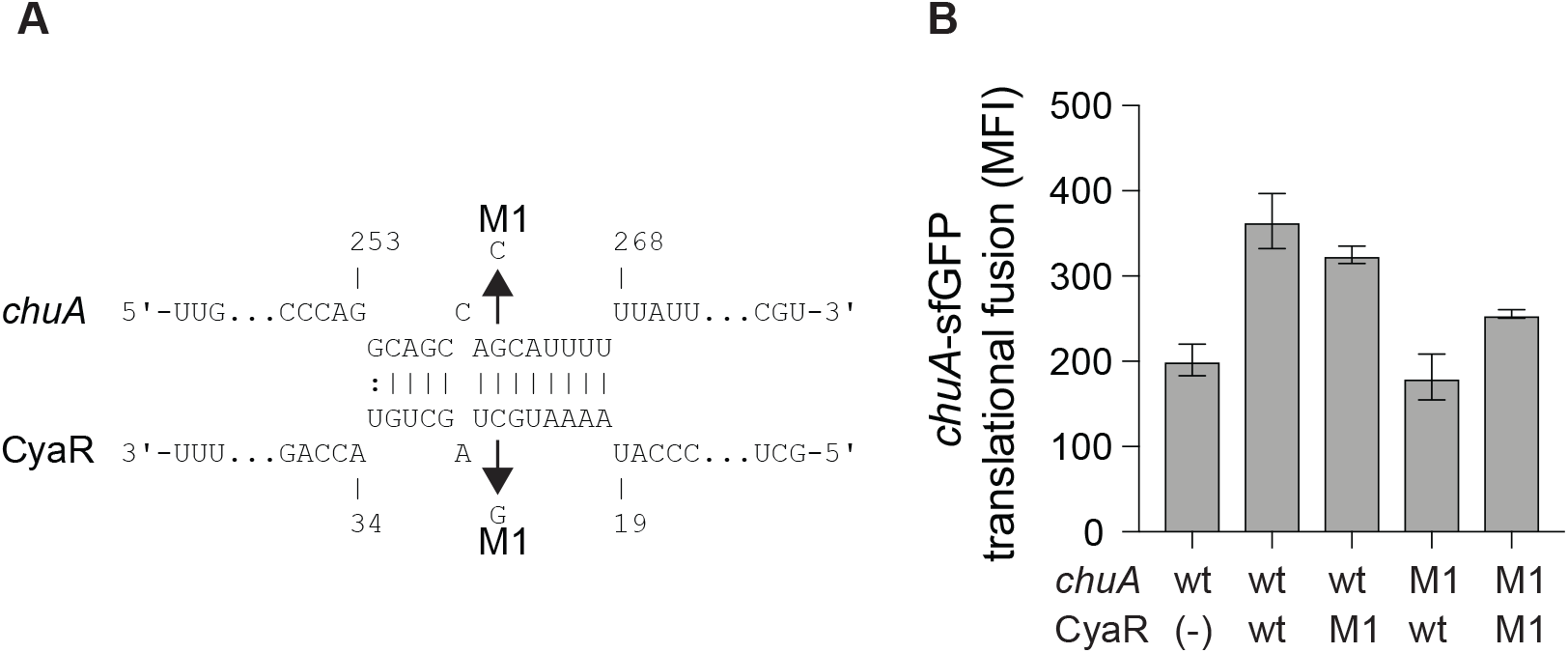
CyaR activates *chuA* translation through direct base-pairing. A. IntaRNA prediction of the *chuA*-CyaR interaction. Compensatory point mutations predicted to disrupt the interaction are indicated by the arrows. B. Fluorescence measurements of wild-type or mutant *chuA*-sfGFP translational fusions in the presence and absence of wild-type or mutant CyaR. Measurements are the mean median fluorescence intensity of three biological replicates (** *p <* 0.01, * *p <* 0.05).

### CyaR activates ChuA translation independently of temperature

Regulation of *chuA* occurs on both the transcriptional and translational level. Transcription of *chuA* is inhibited by Fur when cells are grown in iron-rich conditions [19], while the level of translation is dependent on the environmental temperature. At 25°C, translation is inhibited by the formation of a FourU RNA thermometer that occludes the ribosomal binding site [34]. This inhibitory structure lies 15 nts downstream of the CyaR binding site. To determine whether CyaR-mediated activation of *chuA* translation occurs by inhibiting the formation of this RNA-thermometer, a U273A point mutation known to disrupt the formation of the FourU hairpin loop was made in the 5’UTR of the *chuA*-GFP translational fusion (Figure 3A) [34]. Expression of wild type and mutant *chuA*-sfGFP translational fusions were monitored in the presence and absence of CyaR at 25°C. Disruption of the RNA thermometer via the U273A point mutation resulted in a 2.6-fold increase in fluorescence compared to the wild-type, confirming the FourU temperature-dependent regulation of *chuA* translation. Disruption of this secondary structure however did not affect CyaR-mediated activation of *chuA*, as a 1.6-fold increase in fluorescence was observed when CyaR was provided with the U273A mutation in *chuA* (Figure 3B). This demonstrated that CyaR-mediated activation of *chuA* acts independently of post-transcriptional inhibition through the FourU RNA thermometer structure.

**Figure 3:**
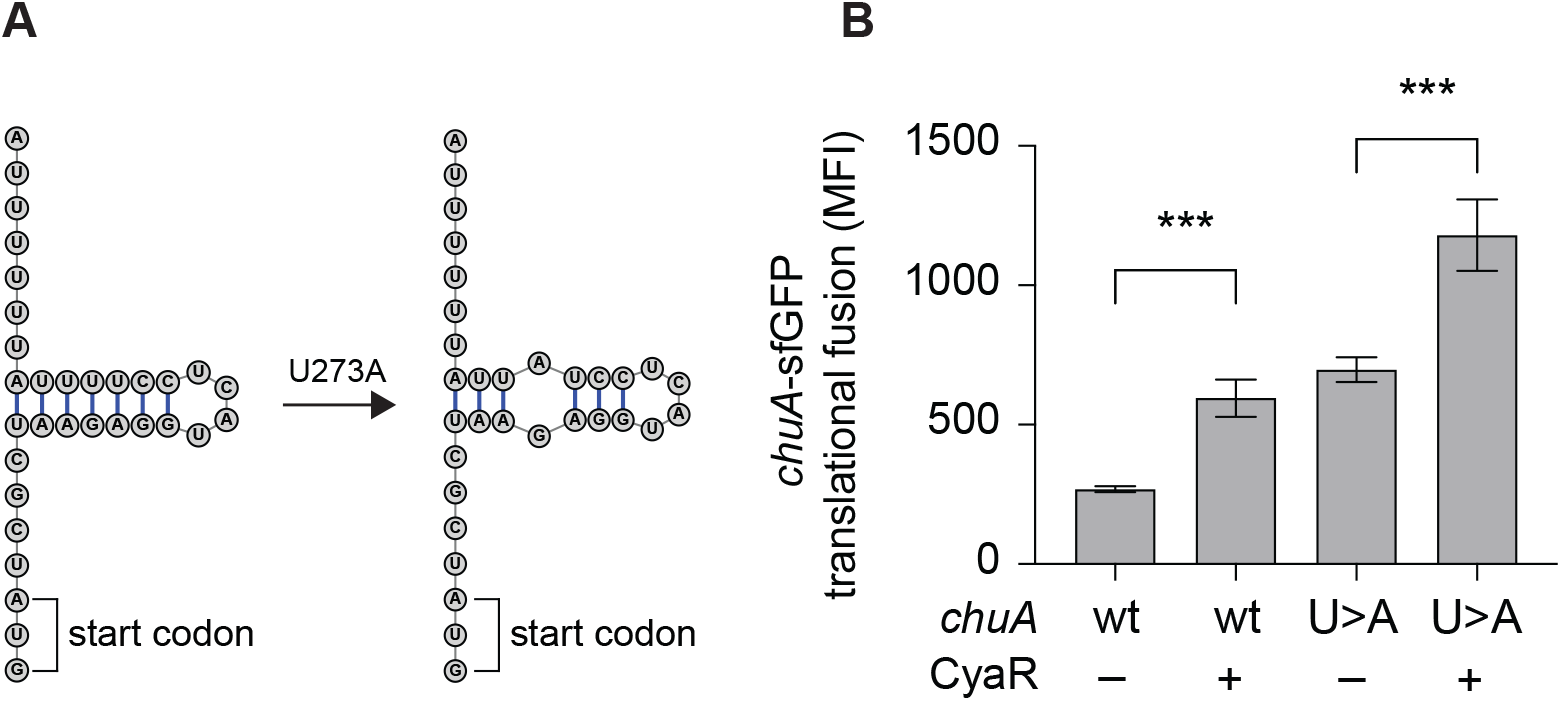
CyaR activates *chuA* translation independently of a FourU RNA-thermometer. A. Diagram indicating the point mutation made to disrupt the formation of the *chuA* FourU RNA thermometer. B. Fluorescence measurements of wild-type and RNA-thermometer disrupted *chuA*-sfGFP translational fusions in the presence and absence of CyaR. Measurements are the mean median fluorescence intensity of three biological replicates. (*** *p <* 0.001).

### The *chuA* 5’UTR contains a sequence that inhibits its expression

While the activation of *chuA* translation by CyaR is not due to disruption of the RNA thermometer, *in silico* folding predictions on the 5’UTR of *chuA* indicated potential secondary structure throughout the 5’UTR, including some that occluded sequences downstream of the ribosomal binding site (Supplementary Figure 2). To understand whether the activating *chuA*-CyaR interaction requires structured sequences within the *chuA* 5’UTR, we progressively removed sequence from the 5’ end in four truncations of the *chuA*-sfGFP translational fusion (Figure 4A). Predicted secondary structure and the presence of Hfq distal face binding motifs ([ARN]_x_) were used as guides in making truncates of the *chuA* 5’UTR [48–50]. A search through the 5’UTR of *chuA* identified ten (ARN)_4_ sites with one mismatch tolerated (ARN_4_m_1_). The first truncate (T1) begins at +57 nt and removes a section that forms two hairpins as well as 9 out of 10 ARN_4_m_1_ sites. The second (T2) and third (T3) begin at +116 and +177, respectively, and each one removes another hairpin from the overall predicted secondary structure while maintaining the RNA-thermometer. The second truncate T2 also removes the last ARN_4_m_1_ motif. The fourth truncate (T4) begins at +217 and leaves only the CyaR binding site, the RNA-thermometer, RBS and the start codon. In the absence of CyaR, the *chuA* 5’UTR T1-T4 truncates caused 2.1-, 1.9-, 2.9- and 9.2-fold increases in *chuA* translation, respectively. CyaR was able to activate truncates T1, T2 and T4 between 20-40% consistent with CyaR activation of the wild type construct (Figure 4B). These results indicate that CyaR does not act through alleviation of an inhibitory secondary structure or a regulatory sequence in the upstream +1 to +217 nt region of the 5’UTR. Notably, removing the region between +177 and +217 nt (T3-T4) of the *chuA* 5’UTR resulted in a dramatic increase in *chuA*-GFP expression (9.2-fold) indicating that the T3-T4 region of the 5’UTR contains sequences or RNA structures that strongly repress expression.

**Figure 4:**
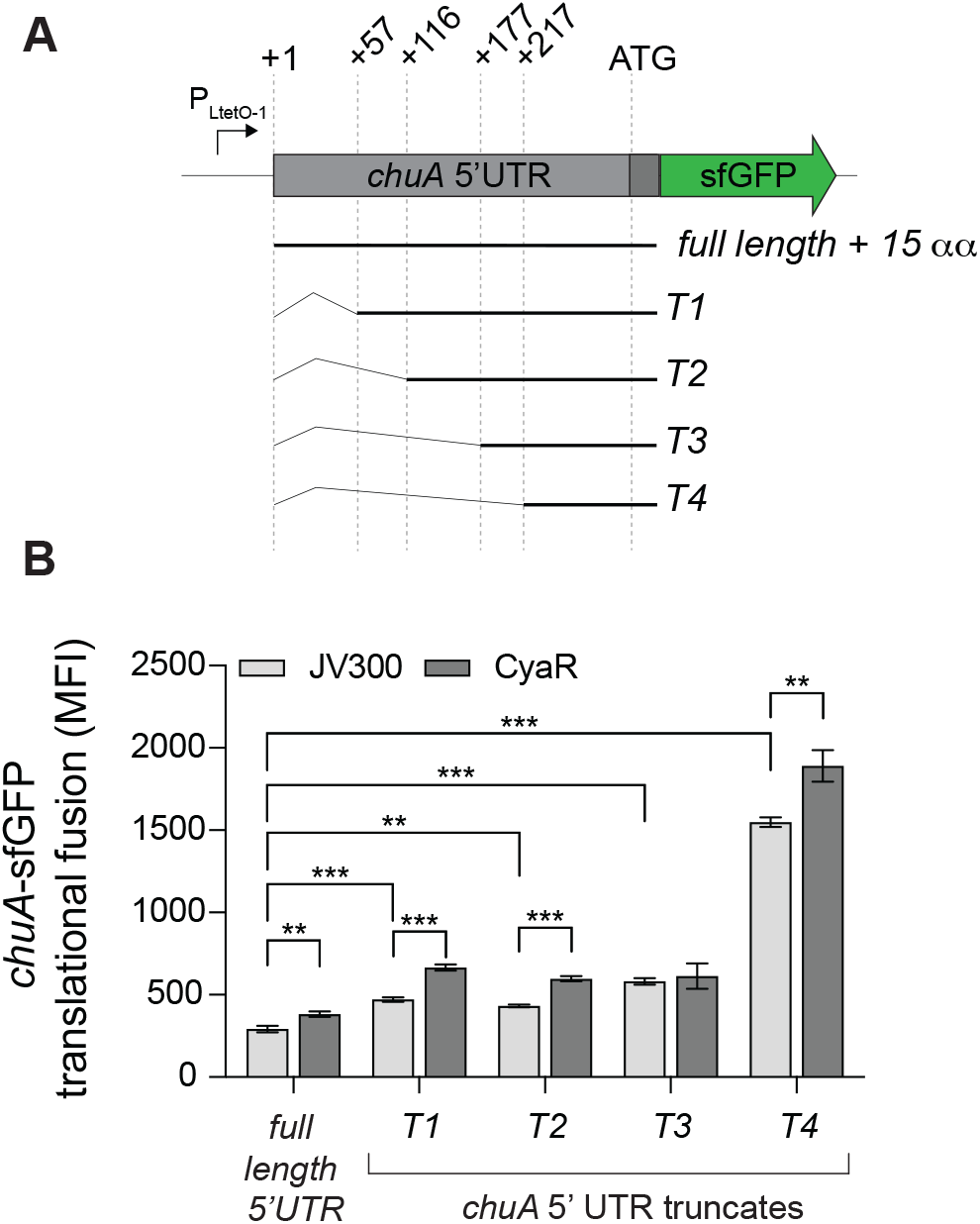
The *chuA* 5’UTR contains a sequence that disrupts its translation independently of CyaR. A. Diagram indicating the positions of the different *chuA*-sfGFP truncates used. B. Fluorescence measurements of full-length and truncated *chuA*-sfGFP constructs in the presence and absence of CyaR. Measurements are the mean median fluorescence intensity of three biological replicates (*** *p <* 0.001, ** *p <* 0.01).

### *chuA* is subject to premature transcription termination by Rho

In *E. coli*, the average 5’ UTR is approximately 25-35 nucleotides in length [51,52]. The 5’UTR of *chuA* is 291 nucleotides, making it an unusually long UTR [34,53]. In commensal *E. coli*, over half of all annotated genes with long 5’UTRs (defined as >80 nts) are prematurely terminated by the transcription termination factor Rho [54]. Further, horizontally transferred genes have been shown to be more susceptible to regulation by Rho-dependent termination [55,56]. By analysing our previously published Term-seq data, we noticed that there was an abundance of 3’ end reads mapped to the 5’UTR of *chuA*, however no Rho-independent terminators were predicted to be in this site using ARNold (Figure 5A) [40,57]. Collectively, these suggested that transcription of the pathogenicity island encoded *chuA* may be prematurely terminated by Rho.

**Figure 5:**
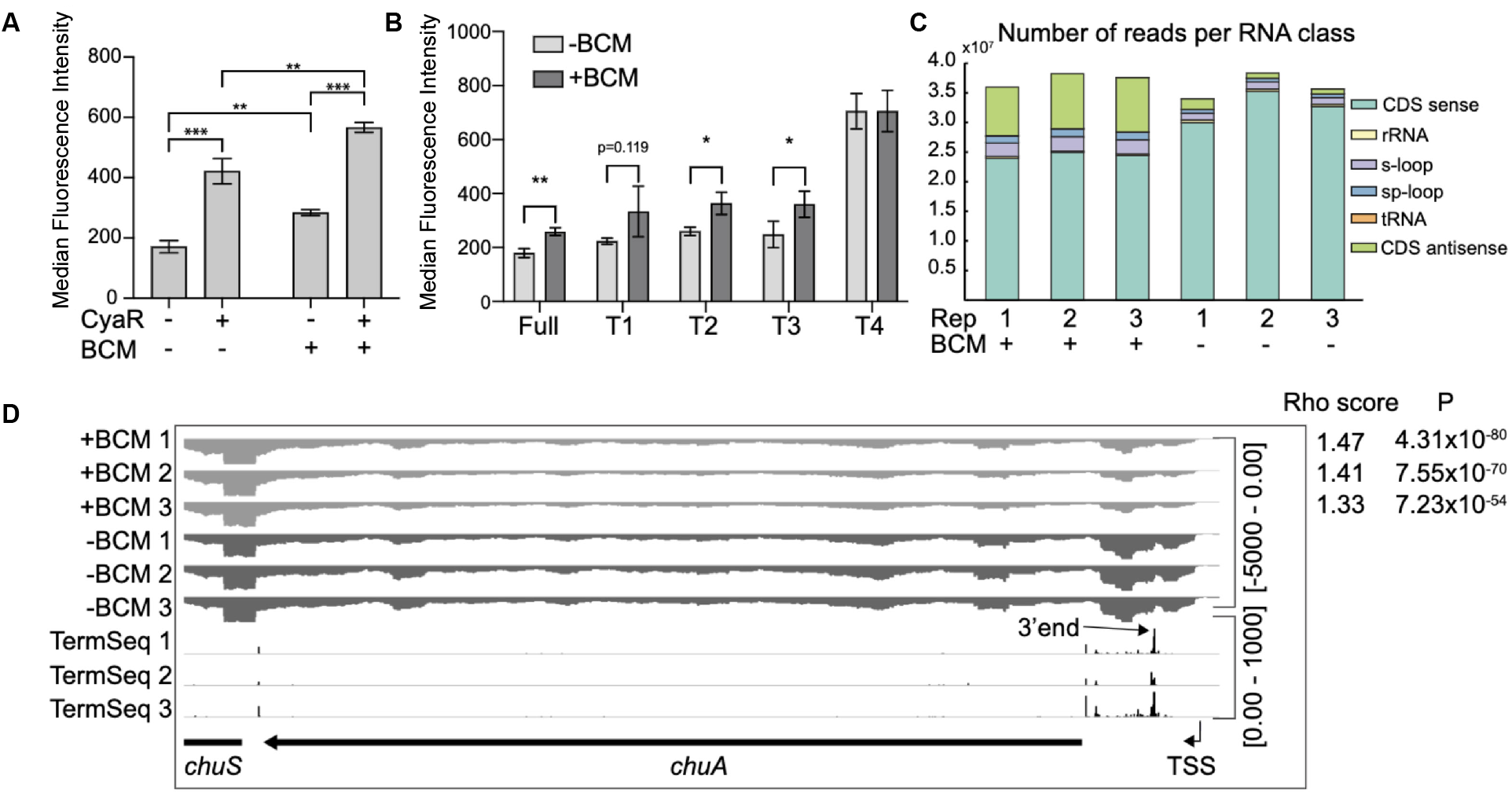
The *chuA* 5’UTR is subject to termination by Rho. A. RNA-seq coverage of *chuA* in EHEC O157:H7 strain Sakai treated and untreated with bicyclomycin. Rho scores for each replicate is shown. TSS is derived from differential RNA-seq data and is indicated by an arrow. Term-seq reads indicating *chuA* 3’ ends are also included. B. Classification and number of reads per RNA class upon treatment of EHEC O157:H7 strain Sakai with bicyclomycin. C. Fluorescence measurements of full-length and truncated *chuA*-sfGFP fusions in the presence and absence of induction with bicyclomycin. D. Fluorescence measurements of *chuA*-sfGFP translational fusion in the presence and absence of CyaR and treatment with bicyclomycin. Measurements are the mean median fluorescence intensity of three biological replicates (*** *p <* 0.001, ** *p <* 0.01, \**p <* 0.05).

To identify Rho-termination sites in EHEC, we sequenced total RNA extracted from cells treated with the Rho-inhibiting antibiotic bicyclomycin [58]. Rho-termination sites were identified by using the approach adopted by [59] that previously identified termination sites in bicyclomycin-treated *E. coli* cells. Consistent with earlier studies we found that horizontally acquired regions and antisense RNAs are enriched among genes that are up-regulated by BCM treatment (Figure 5B) [56,60]. Read-through scores were calculated to assess Rho-dependant termination using previously published scripts [59]. Read-through transcription of the *chuAS* transcript was up-regulated at least 1.35 fold by BCM treatment indicating that Rho prematurely terminates transcription of *chuA* (Figure 5A). To verify that *chuA* was subject to premature transcription termination by Rho, we measured the fluorescence of inducible full-length and truncated *chuA*- GFP translational fusions in the presence or absence of BCM (Figure 5A). Treatment with BCM resulted in a ∼1.4-fold increase in fluorescence for the full-length and T1-T3 *chuA*-GFP constructs. No significant change was observed for *chuA* truncate T4 (Figure 5C). Taken together, these results indicate that *chuA* is negatively regulated by Rho terminator and this repression requires the 30 nt between +177 and +217 nt of the *chuA* 5’ UTR.

Small RNAs can regulate Rho-dependant termination by altering the accessibility of Rho utilisation sites, either as a by-product of translation inhibition, or by directly binding to the *rut* site itself [54,61–63]. We hypothesised that CyaR may activate *chuA* translation by preventing premature Rho termination within the 5’UTR. To test this, we measured the fluorescence of our *chuA*-GFP translational fusion in the presence or absence of BCM and CyaR. CyaR was able to activate translation of *chuA* in the presence of BCM, indicating that CyaR activates *chuA* independently of Rho termination (Figure 5D).

Collectively our data demonstrate that *chuA* expression is independently regulated by CyaR sRNA, Rho terminator, and the FourU RNA thermometer. With previously published data demonstrating regulation by Fur, RyhB, and Esr41 [19,36] it appears that *chuA* expression is controlled by eight regulatory inputs that control expression through six different transcriptional and post-transcriptional mechanisms.

## DISCUSSION

To successfully grow and colonise a host, pathogens utilise systems that allow them to retrieve trace minerals required for growth that are normally sequestered. Various pathotypes of *E. coli*, including EHEC and UPEC, utilize the ChuA outer membrane haem receptor to transport host derived haem. Transcription of *chuA* is repressed in the presence of iron by Fur [19], while translation is regulated by temperature through a FourU RNA-thermometer that occludes the ribosomal binding site [34]. The combination of these two modes of regulation allows the pathogen to sense two signals associated with the host environment (low iron and high temperature) and activate expression of the haem receptor inside the host. Notably, each signal in isolation would be generate many false positives when deciding on whether the cell had entered a mammalian host.

The natural order imposed by transcription and post-transcriptional regulation creates an AND-logic gate [84], where low iron levels AND high temperature are required for expression of *chuA* in the host. Here we have shown that *chuA* is controlled by two additional post-transcriptional signals: repression through Rho termination and activation through CyaR. Importantly, CyaR acts independantly of both Rho and the FourU thermometer creating a post-transcriptional OR-logic gate. The effect of the FourU thermometer and CyaR are additive, and either regulator can activate independently of the other. In Boolean terms it appears that *chuA* uses an AND-OR logic gate where expression requires low iron AND (high temperature OR CyaR).

Transcription of CyaR is activated by the global regulator Crp when cyclic-AMP levels are high, such as in the absence of glucose [64]. EHEC colonise the colon where the primary source of carbon is mucins. *Bacteroides thetaiotaomicron (Bt)* can cleave mucins to release sugars that are utilized by EHEC [65,66]. During colonization of the gastrointestinal mucus layer and epithelium, EHEC is likely to encounter varying niches that include both glycolytic and gluconeogenic environments [67]. Sensing variation in the availability of sugars and oxygen availability have been suggested to be a key regulatory signals that allow EHEC to determine location in the gastrointestinal tract and proximity to the epithelium [68]. Close to the intestinal epithelium, the relative absence of microorganisms that metabolise and release sugars from mucins creates at gluconeogenic environment [69]. Lower levels of fucose, and higher levels of succinate reduces the level of the fucose-sensing two-component system FusKR that induces expression of Cra. In gluconeogenic environments, Cra can enhance binding of the cAMP receptor protein Crp to its targets [70]. Activation of Cra can induce expression of genes required for T3SS and adhesion to the epithelium [68,71,72]. Expression of Crp can also be activated in the presence of Cra [73]. We propose that the gluconeogenic environment encountered at the gastrointestinal tract provides an activating signal for CyaR that has been incorporated into *chuA* regulation as an additional signal that indicates host colonization and haem availability.

The mechanism of CyaR activation of *chuA* expression remains unclear although we have ruled out three probable mechanisms: CyaR acts through direct base-pairing with the *chuA* 5’UTR rather than general titration of Hfq, CyaR does not disrupt the FourU RNA thermometer to override temperature-dependance, and CyaR does not inhibit premature Rho termination of *chuA*. While these results have uncovered two additional post-transcriptional regulators of *chuA* (CyaR and Rho) they also suggest that there exists an additional repressive element in the *chuA* 5’UTR that is overcome by CyaR binding.

Using truncates of the *chuA* 5’UTR we have found that sequence between +177 and +217 nt (T3 and T4) represses *chuA* expression almost 10-fold. These results correlate with the region required for premature Rho termination and the presence of a predicted Rho utilization and RNAP pause site. We suggest that Rho associates with the nascent *chuA* transcript between +177 and +217 nt to terminate transcription. Small RNAs can promote or occlude *rut* sites to control Rho termination in response to regulatory signals [54,62]and it remains plausible that Rho termination is modulated by yet another regulatory signal.

The *chuA* encoded haem receptor is transcribed as a bicistronic transcript with the haem oxygenase *chuS*. In previous work we showed that *chuS* expression is repressed by the sRNA FnrS that is induced under anaerobic conditions, consistent with ChuS requiring oxygen for activity. FnrS repression is over-ridden by the sRNA sponge AsxR transcribed from the Shiga toxin 2 encoding bacteriophage Sp5 [35]. Collectively, expression of *chuAS* is subject to an impressive level of post-transcriptional regulation that provides complex integration of environmental signals beyond low iron (transcriptional regulation by Fur). These post-transcriptional signals appear to provide a much more accurate interpretation of the environment to indicate whether haem transport is required (ie: whether EHEC is at the gastrointestinal epithelium). The logic of the regulatory circuit that controls *chuAS* expression is outlined in Figure 6.

**Figure 6:**
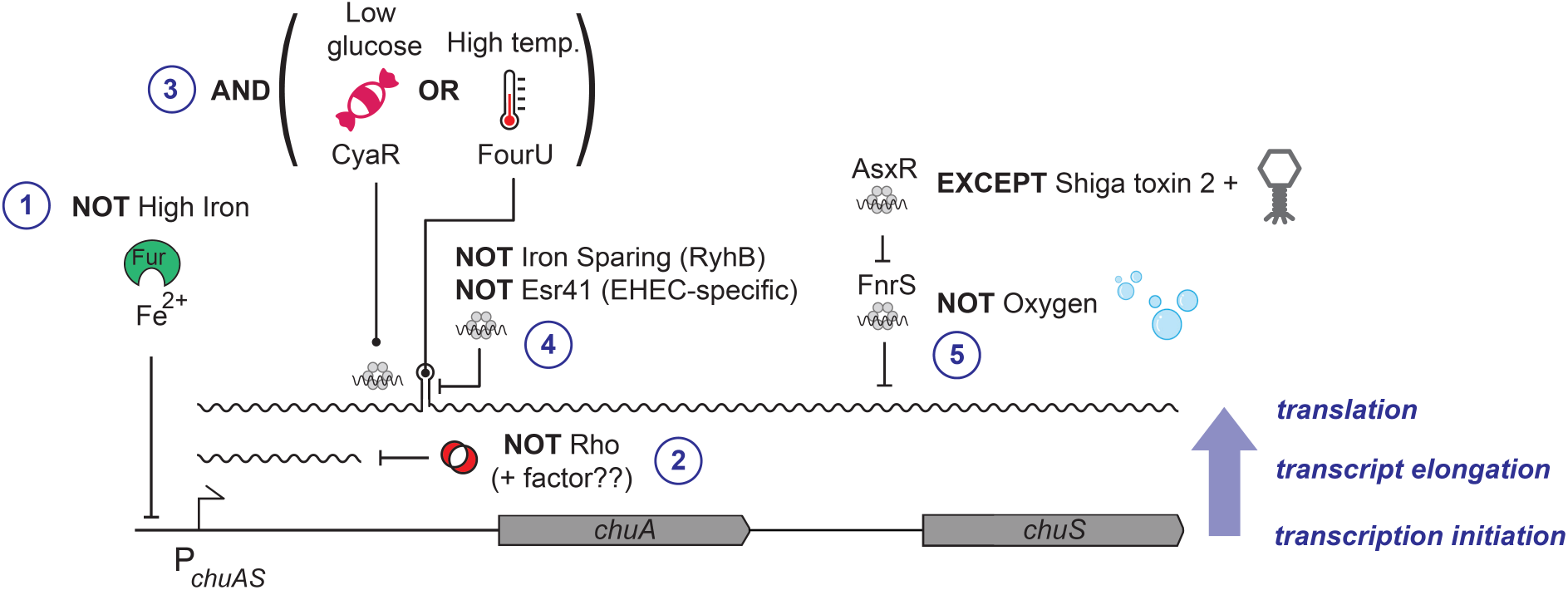
Model of *chuAS* regulation. Regulation of *chuAS* expression occurs at the transcription intitation, transcript elongation, and translational levels (indicated blue, right). The dependance of each layer on the previous creates conditional regulatory AND gates [84]. (1) Repression of *chuAS* transcription by Fe-loaded Fur is relieved under iron-limited conditions allowing transcription initiation. (2) Elongating RNAP is prematurely terminated by Rho within the 5’UTR of *chuA*. An additional factor (potentially another sRNA) is likely to modulate Rho termination and allow read-through transcription. (3) Translation of the *chuAS* transcript is activated by the cAMP-responsive sRNA CyaR or temperatures ≥ 37°C that melt the FourU thermometre. (4) Melting of the *chuAS* mRNA FourU element can be overridden under iron-sparing conditions by the sRNA RyhB or the EHEC-specific sRNA Esr41 that repress translation from the exposed RBS. (5) The downstream haem oxygenase encoded by *chuS* is repressed under aerobic conditions by the sRNA FnrS, except in EHEC where repression is relieved by the sRNA sponge AsxR carried by the Shiga toxin 2-encoding bacteriophage.

Post-transcriptional regulation appears ideally suited to this layering of logic gates because the natural order (dependance) imposed by transcriptional and post-transcriptional signals creates AND gates [84]. There are at least three ordered layers of AND gates in the *chuA* transcript that need to be satisfied before the next layer of signals are incorporated. Fur-dependant transcription, Rho-dependent termination, and post-transcriptional control through sRNAs or the FourU thermometer. Together these form an elegant set of AND, OR, and NOT gates that interpret the environment and genetic background of the host (the later through the EHEC-specific sRNAs AsxR and Esr41)(Figure 6). The extensive post-transcriptional logic of the *chuAS* operon suggests that the commonly quoted purpose of sRNAs for “fine-tuning” expression over-simplifies the opportunities that layered post-transcriptional regulation presents for creating complex regulatory logic gates to integrate and interpret environmental signals.

## MATERIALS AND METHODS

### Bacterial strains and growth conditions

Bacterial strains, oligonucleotides and plasmids used for this study are listed in Supplementary table 1. *E. coli* was routinely grown at 37°C in liquid Luria-Bertani (LB) broth or on solid LB-agar plates. Bacterial media was supplemented with ampicillin (100 μg/mL) or chloramphenicol (34 μg/mL) where appropriate. For inhibition of Rho termination, bicyclomycin (50 μg/mL) was added to exponential phase (OD600 = 0.6) cultures for 30 minutes before being assayed.

### In silico prediction of interacting sRNAs

A list of published sRNAs present in enterohaemorrhagic *E. coli* was taken from the Bacterial Small Regulatory RNA Database [44]. sRNAs listed as not being Hfq-binding were filtered out, and the sequences of the remaining sRNAs were input into IntaRNA [42,43,74] to search for interactions with the 5’UTR of *chuA*. sRNAs that were predicted to bind to regions of the *chuA* 5’UTR that were not Hfq-binding were removed from consideration.

### Construction of GFP-translational fusions and sRNA expression vectors for testing sRNA-mRNA interactions

Plasmids pXG10SF containing the full-length and truncated *chuA*-GFP translational fusion and pZE12 carrying candidate sRNAs were cloned according to method described in [75]. Primers used for cloning are listed in Supplementary Table 1.

Mutations were made in the *chuA* or sRNA sequences by using the Quikchange XL mutagenesis kit (Agilent) according to the manufacturer’s instructions. Primers for mutagenesis were designed using the Quikchange primer design program (https://www.agilent.com/store/primerDesignProgram.jsp).

### Confirmation of sRNA-*chuA* interactions using the sfGFP 2-plasmid system

The expression of sfGFP was monitored and quantified with and without candidate sRNAs using a BD FACSCanto II or a BD LSRFortessa™ Special Order Research Product cell analyser. Fluorescence was measured using a 530/30 nm bandpass filter. FSC and SSC were also measured to gate the bacterial population. For each sample, at least 100,000 events were recorded. Data was analysed using FlowJo software (BD) and statistics were calculated using Prism 8 (GraphPad) to obtain each sample’s mean median fluorescence intensity (mean MFI). *p*-values were calculated using a standard two-tailed student’s t-test.

Plasmids expressing the wild type or mutant *chuA*-GFP translational fusion and those expressing candidate sRNAs were co-transformed into *E. coli* DH5α or TOP10F’. Three colonies from each transformation were purified and grown overnight in LB broth. Overnight cultures are subcultured 1/100 in 0.22 μm filtered LB broth and grown to exponential phase (OD_600_ = 0.6). Expression of the *chuA*-GFP translational fusions were induced in TOP10F’ using 200 nM of anhydrotetracycline. These are diluted five-fold in 0.22 μm-filtered PBS, then read on the flow cytometer as described above. Plasmid pJV300 expressing a scrambled sRNA, and pXG1 and pXG0 plasmids expressing GFP and Lux, respectively, were used as controls.

### RNA secondary structure prediction

The secondary structure for the 5’UTR of *chuA* was predicted using the mfold (unafold.rna.albany.edu/?q=mfold) and RNAfold webservers [76,77]. Figures were drawn on RNAStructure version 6.0.1 [78]. ARN motifs were detected using custom scripts previously used in [35].

### RNA sequencing of bicylomycin-treated cells

Three single colonies of *E. coli* O157:H7 str. Sakai *stx*- were grown for 16 hours in LB broth. The following day, these subcultured 1/100 in MEM-HEPES supplemented with 0.1% glucose and 250 nM Fe(NO_3_)_3_. At an OD_600_ of 0.75, cultures were split into two (treated and untreated), and 50 μg/mL of bicyclomycin was added to the treated sample. Cultures were incubated for a further 15 minutes, followed by addition of RNA stop solution (5% water-saturated phenol in ethanol). RNA was extracted using guanidinium thiocyanate-phenol as previously described in [79]. Genomic DNA was digested using RQ1 DNase (Promega) and RNA was cleaned using another phenol-chloroform extraction. Total RNA was ribodepleted using the Zymo-Seq RiboFree Total RNA library kit and libraries were prepared using the Illumina NextSeq 500/550 Mid-Output Kit. Samples were 2×75 bp paired-end sequenced on an Illumina Nextseq 500 platform. Library preparation and sequencing were performed by the Ramaciotti Centre for Genomics in the University of New South Wales, Sydney, Australia.

### Identification of Rho-terminated transcripts

Alignment and DEseq analysis of RNA sequencing output was done using the “align”, “coverage”, “gene-quanti” and “deseq” modules of READemption v1.0 [80] using default settings.

To identify Rho-terminated transcripts, fastq files were re-aligned to the *E. coli* O157:H7 str. Sakai genome (accession number: NC_002695.2) using BWA-MEM (v0.7.17) [81]. BAM files were generated using samtools (v.1.10) [82] and 3’ end read counts were calculated using the genomecov tool from bedtools (v2.27.1) [83]. Rho readthrough scores were calculate using the program RhoTerm-Peaks using a window size of 250 nt [59].

## FIGURE LEGENDS

**Supplementary Figure 1:**
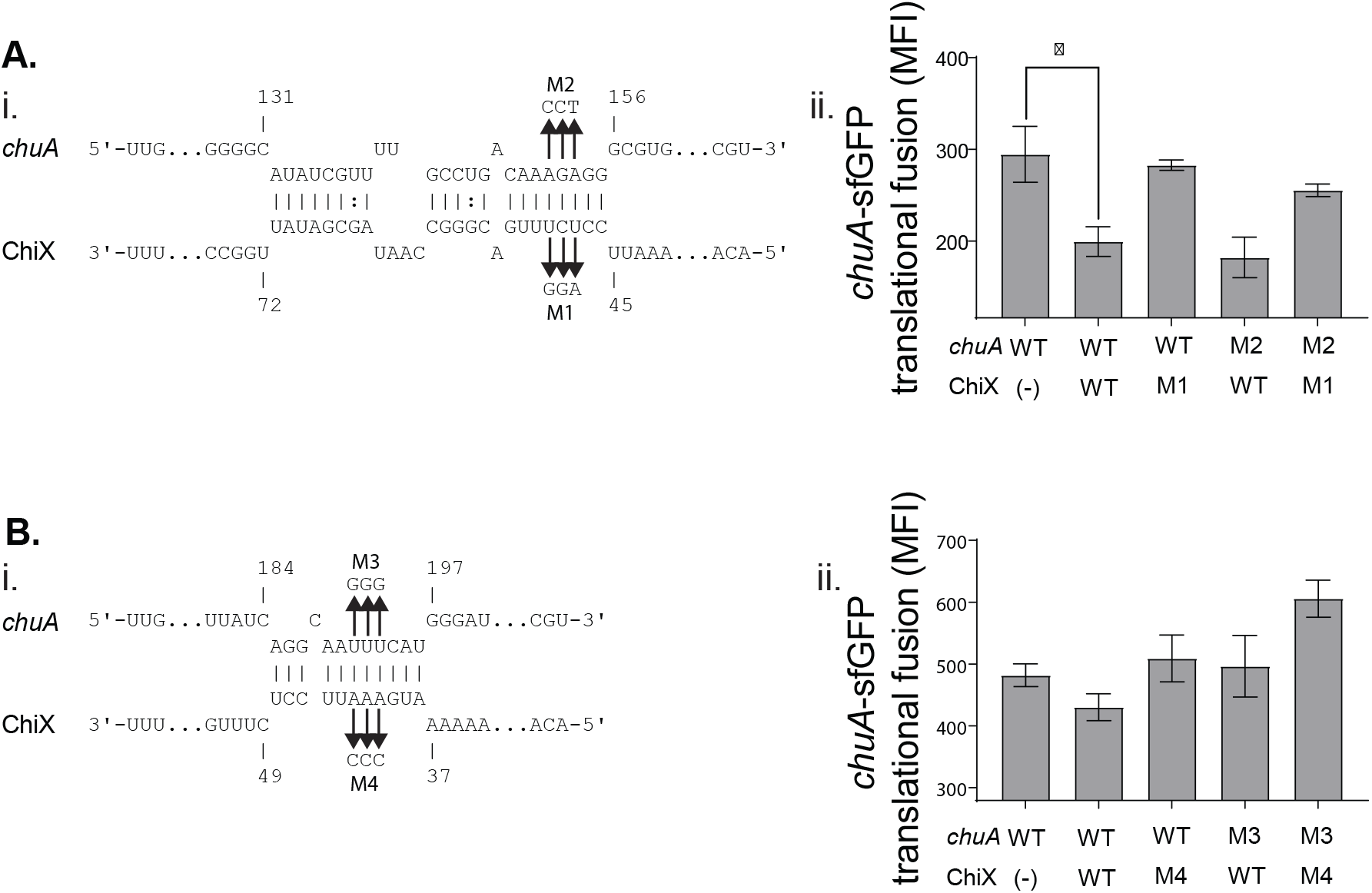
ChiX indirectly represses *chuA* translation. A-B. (Left) IntaRNA prediction of the *chuA*-ChiX interaction. Compensatory point mutations predicted to disrupt the interaction are indicated by the arrows. (Right) Fluorescence measurements of wild-type or mutant *chuA*-sfGFP translational fusions in the presence and absence of wild-type or mutant ChiX. Measurements are the mean median fluorescence intensity of three biological replicates.

**Supplementary Figure 2:**
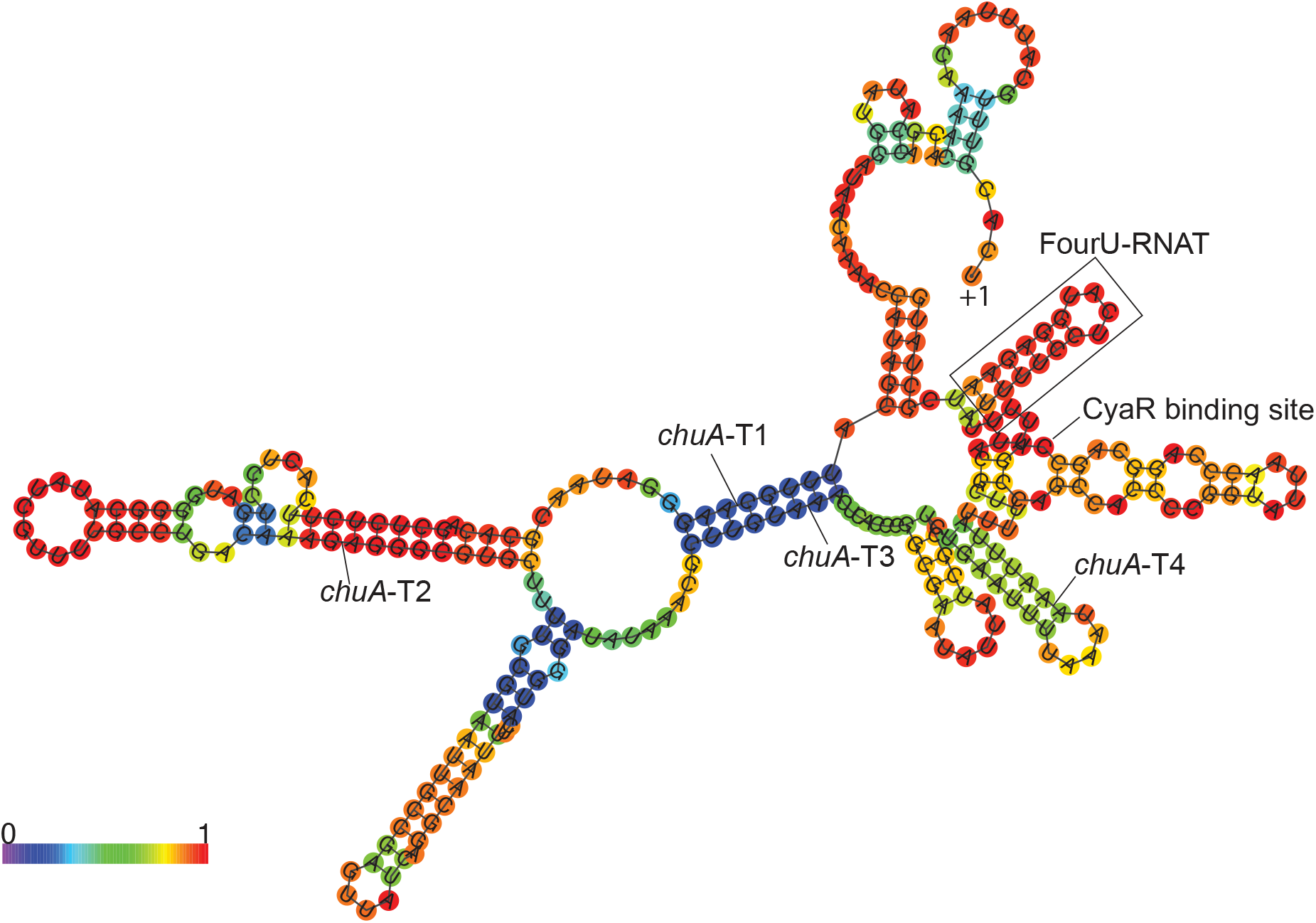
Secondary structure of the *chuA* 5’UTR as predicted by RNAfold. The +1 site, CyaR binding site, FourU RNA thermometer and sites where truncations were made are indicated.

**Supplementary Table 1**: Primers, bacterial strains and plasmids used in this work.

